# ChromBERT-tools: A versatile toolkit for context-specific embedding of transcription regulators across different cell types

**DOI:** 10.64898/2026.02.04.703739

**Authors:** Qianqian Chen, Zhaowei Yu, Yong Zhang

## Abstract

**Motivation:** Representations that capture the genome-wide context of transcription regulators are critical for establishing a shared backbone for flexible transcription modeling and in silico regulatory analysis. Yet, current embeddings predominantly rely on limited modalities, such as gene co-expression or static protein features, offering an incomplete perspective that ignores context-dependent transcription regulator activities across the genome. The lack of transcription regulation-informed embeddings, paired with the absence of a user-friendly and lightweight toolkit for their generation, adaption to different cell types and interpretation, impedes the capture of the regulatory logic that underpin cellular states and functions.

**Results:** To address this need, we present ChromBERT-tools, a lightweight toolkit designed to operationalize regulation-informed embeddings derived from a foundation model pre-trained on the comprehensive landscapes of human and mouse transcription regulators. ChromBERT-tools provides user-friendly command-line interfaces (CLIs) and Python APIs to achieve two primary goals: (i) generating cell-type-agnostic embeddings that capture the semantic representations of individual regulators and their combinatorial interactions, serving as biological priors of transcription regulator modality to enhance transcription regulation modeling and rule interpretation; and (ii) generating cell-type-specific embeddings via fine-tuned model variants, which support in silico inference of regulatory roles of transcription regulators in cell types with scarce experimental data. The toolkit streamlines end-to-end workflows for embedding generation, adaption to different cell types and interpretation towards biological inferences such as regulator-regulator interaction across the genome and key regulators determining cell identity or cell state transition.

**Availability and implementation:** ChromBERT-tools is freely available at https://github.com/TongjiZhanglab/ChromBERT-tools, with documentation at https://chrombert-tools.readthedocs.io/en/latest/.

## 1 Introduction

Representation learning has emerged as a transformative framework for decoding the complexity of biological data. By projecting high-dimensional data, such as transcriptomes, proteomes and genomes, into compact embedding spaces, this approach reveals latent semantic patters that facilitate the discovery of novel biological logic. Crucially, these context-aware representations provide a versatile foundation for transfer learning, enabling the robust adaptation of general biological insights to diverse, data-scarce downstream applications (Theodoris *et al*, 2023). Recently, the power of this paradigm is unlocked by the large-scale pre-training of deep language models, which learns the fundamental ‘grammar’ of biological sequences from vast unlabeled corpora. Built on Transformer architecture, these models employ attention mechanism to master long-range dependencies (Devlin *et al*, 2018; Theodoris *et al*, 2023; Vaswani *et al*, 2017), ensuring that the resulting representations are not only biologically rich but also highly transferable(Fu *et al*, 2025). Furthermore, this foundation support multimodal integration, aligning disparate representation spaces to enable seamless cross-modal retrieval and holistic system-level modeling (Hingerl *et al*, 2025; Khan *et al*, 2025).

Transcription regulators (TRs) serve as master regulators of cellular identity, orchestrating dynamic regulatory landscapes through precise, combinatorial interactions at *cis*-regulatory elements (CREs) (Kim and Wysocka, 2023; Stampfel *et al*, 2015). Recent transformer-based models have successfully captured distinct dimensions of these regulators: single-cell foundation models, such as scGPT, excel at mapping cell-type-specific gene-gene interaction networks from expression data (Cui *et al*, 2024), while protein language models, such as ESM-2, effectively encode intrinsic biochemical and structural potentials from amino acid sequences (Lin *et al*, 2023). Nevertheless, these approaches remain disconnected from the genome. By relying on static sequences or co-expression correlations, they fail to explicitly model the *trans*-acting mechanisms of TFs, the direct genome-wide binding landscapes and combinatorial co-occupancy patterns required to understand cooperative regulation. To bridge this gap, we pre-trained ChromBERT on large-scale TR ChIP-seq data, demonstrating that regulation-informed representations can effectively capture these missing physical mechanisms and generalize across cell types (Yu *et al*, 2026). Unlike sequence- or expression-derived baselines, its regulation-informed embeddings explicitly encode context-specific TR-chromatin associations and combinatorial TR-TR co-occupancy, thereby resolving the disconnect between computational models and the dynamic genome context.

ChromBERT has demonstrated significant efficacy in modeling genomic regulation through large-scale pre-training. Translating these high-dimensional representations into practical insights often involve tailoring the model to specific biological questions, from static transcription regulator crosstalk to dynamic cell context-aware regulatory architecture. To facilitate this and maximize the model’s general-purpose utility, we introduce ChromBERT-tools, a comprehensive suite of utilities designed to bridge the gap between raw input interested genomic regions or regulators and model inference. Rather than requiring users to build custom pipelines, ChromBERT-tools provides a standardized, end-to-end environment for leveraging ChromBERT’s capabilities. Through flexible CLIs and Python APIs, the toolkit manages input standardization and embedding generation, including cell-type-agnostic and cell-type-specific regulator embeddings, and biologically inference from embeddings, enabling users to seamlessly apply the model to complex tasks and interpret biological insights, with efficiency and precision.

## 2 Toolkit description

### 2.1 Generation of regulation-informed embeddings

ChromBERT-tools implements an end-to-end framework for generating regulation-informed embeddings through two distinct workflows (Fig. 1A): a cell-type-agnostic workflow that produces pre-trained embeddings, and a cell-type-specific workflow that fine-tunes ChromBERT to the provided cellular context. These workflows are accessible via CLIs for reproducible execution and Python APIs for integration into custom analysis pipelines (Table S1).

**Fig. 1.**
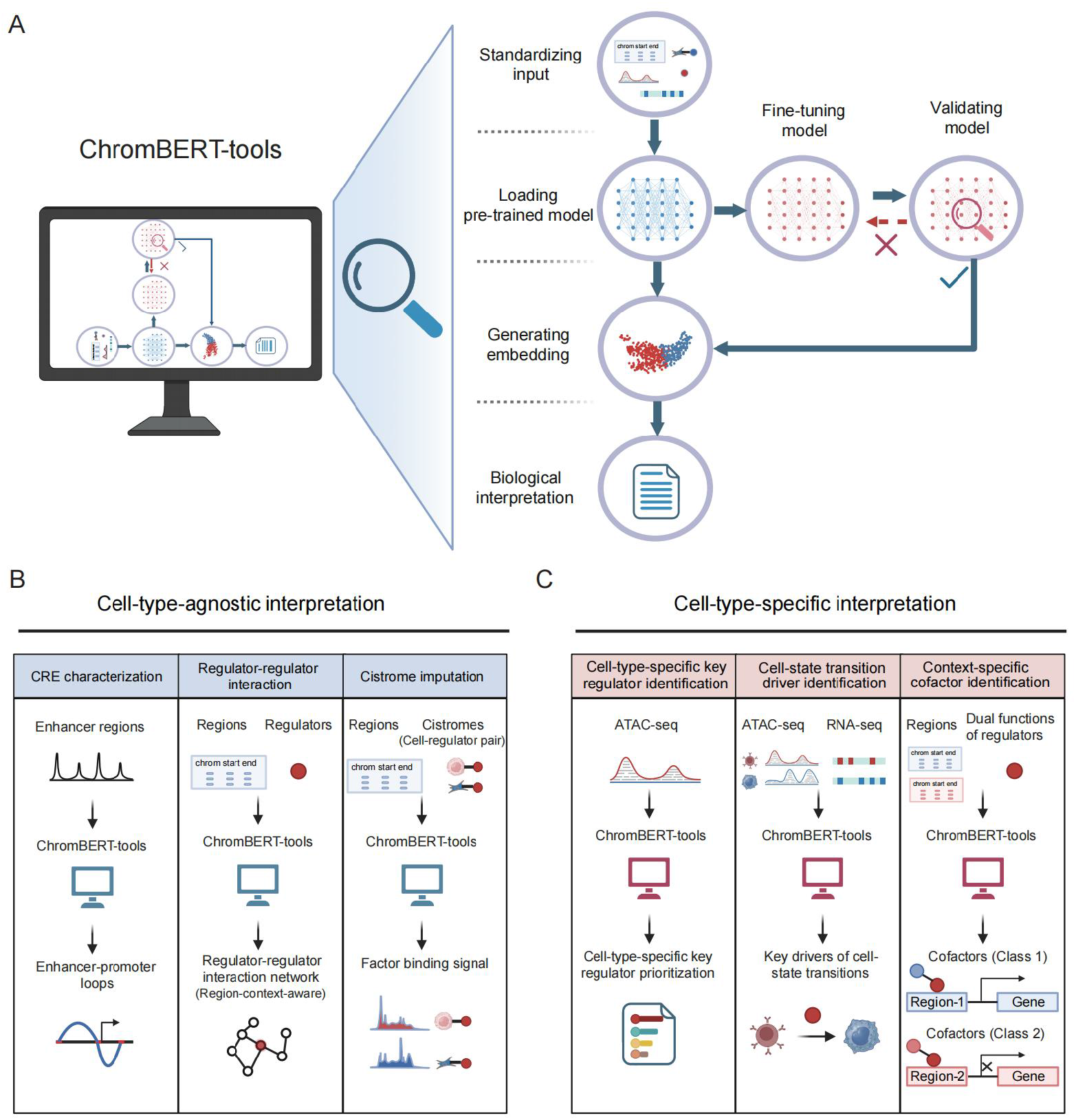
Workflow of ChromBERT-tools. (A) An end-to-end workflow to generate pre-trained or fine-tuned embeddings for biological interpretation. The workflow includes: converting raw data into model-compatible inputs, loading a pre-trained ChromBERT checkpoint, optional fine-tuning with model validation, embedding generation for regulators and regions, and downstream analyses to derive biological insights. (B) Cell-type-agnostic interpretation. Using embeddings from the pre-trained model, ChromBERT-tools supports three representative analyses from left to right: (i) CRE characterization by region-embedding similarity to identify functional relationships among input CREs, such as enhancer-promoter loops; (ii) region-context-aware regulator-regulator interaction inference via regulator-embedding similarity to capture co-regulation relationships for user-specified regions and regulators; and (iii) cistrome imputation to predict TR binding signals over specified regions for user-specified cell-regulator pairs. (C) Cell-type-specific interpretation. Using embeddings generated by fine-tuned model variants, ChromBERT-tools supports three representative analyses from left to right: (i) cell-type-specific key regulator prioritization using regulator embeddings from a model fine-tuned on cell-type ATAC-seq; (ii) cell-state transition driver identification using regulator embeddings from models fine-tuned on paired start and end states with ATAC-seq and/or RNA-seq; and (iii) context-specific cofactor identification using regulator embeddings from a fine-tuned model under distinct region-set contexts to reveal context-dependent cofactor dependencies.

To support diverse experimental systems, ChromBERT-tools includes built-in configurations for human (hg38) and mouse (mm10) reference assemblies. Users define the genome assembly (--genome) and the desired genomic resolution (--resolution), with supported bins ranging from 200 bp to 4 kb for human, and 1 kb for mouse.

#### 2.1.1 Standardizing input

ChromBERT-tools generates embeddings for approximately 1,000 regulators at genomic regions of interest. The toolkit accepts regions in BED format, mapping them to the model’s reference bins at the user-specified resolution. To enable the fine-tuning required for cell-type-specific embeddings, the tool supports user-provided BigWig tracks (e.g., ATAC-seq/DNase-seq). It summarizes these accessibility signals over the input regions to serve as objectives for model adaptation. Further details are provided in the Data processing section of the Supplementary Methods.

#### 2.1.2 Loading pre-trained model

ChromBERT-tools streamlines the initialization process by automatically selecting and loading the appropriate pre-trained ChromBERT checkpoint matching the specified reference genome and resolution. It simultaneously manages all required auxiliary resources, such as reference files and metadata, ensuring a seamless setup for downstream tasks.

#### 2.1.3 Fine-tuning model

For cell-type-specific embedding generation and downstream interpretation tasks such as cell-state transition driver prioritization that require fine-tuning, ChromBERT-tools automatically selects the appropriate training loss functions and supervised metrics. Details are provided in the Model fine-tuning section of Supplementary Methods.

To balance computational efficiency with accuracy, the toolkit offers two fine-tuning modes: (i) Fast mode: Designed for rapid iteration, this mode downsamples the input to a maximum of 20,000 regions. This substantially reduces runtime (typically completing within tens of minutes on an NVIDIA A100 GPU) while maintaining performance comparable to full training (Figs. S2–S4); (ii) Full mode: Utilizes all available training regions to maximize accuracy when runtime constraints permit.

#### 2.1.4 Validating model

To ensure the model is robustly adapted to the target regulatory context, the fine-tuned checkpoint undergoes automated quality control. Recognizing that fine-tuning on high-variance biological data can occasionally yield suboptimal convergence, ChromBERT-tools evaluates the model against reliability criteria and automatically restarts training if these metrics are not met. This ensures that downstream analyses are based on stable, high-quality representations. Details are provided in the Model validation section of the Supplementary Methods.

#### 2.1.5 Generating embeddings

ChromBERT-tools computes regulation-informed embeddings for each input genomic region at two levels: (i) Regulator-level: The model outputs a 768-dimensional vector for every regulator at the given genomic region (1,073 regulators for human; 703 for mouse); (ii) Region-level: Optionally, users can derive a summarized embedding for a genomic region by performing mean-pooling across all regulator vectors for that region.

### 2.2 Biological interpretation of regulation-informed embeddings

ChromBERT-tools interprets regulation-informed embeddings to infer regulatory architecture using two primary metrics. First, it computes pairwise cosine similarity of embeddings to assess relationships between entities. High similarity between regulator embeddings indicates consistent co-binding patterns, suggesting shared regulatory programs or functional synergy (e.g., participation in overlapping pathways). Likewise, high similarity between region embeddings implies functional linkage or shared regulatory properties (e.g., classifying both as promoters or enhancers). Second, the tool quantifies embedding shift (defined as 1 – cosine similarity) to measure how a specific regulator changes across contrasting contexts (e.g., differential accessible regions versus unchanged regions during a cell-state transition). Larger shifts reflect stronger context-dependent alterations in a regulator’s role and are utilized to prioritize key regulators driving regulatory dynamics. Further details are provided in the Embedding analysis section of the Supplementary Methods.

Below, we highlight representative biological interpretation using these metrics, based on the human hg38 genome at 1 kb resolution. For each application, ChromBERT-tools provides the corresponding CLIs and APIs (Fig 1B, C, Table S1).

#### 2.2.1 Context-aware regulator-regulator interaction

Characterizing co-binding partnerships and cofactors dependencies helps to interpret combinatorial regulation. ChromBERT-tools addresses this by generating cell-type-agnostic, context-aware embeddings for each regulator. For input regions of interest, ChromBERT calculate the pairwise similarity of regulator embeddings at the given regions as edge weights to construct regulator-regulator networks reflecting their co-binding and co-regulatory relationships. ChromBERT-tools provides the “infer_regulator_network” subcommand to implement this workflow end-to-end.

We demonstrated the utility of this sub-command by deriving a regulator-regulator network based on CTCF peaks on chromosome 1, focusing on the CTCF-centered subnetwork for illustration (Supplementary Methods). The inferred network successfully recovered the canonical CTCF-cohesin module, a conserved complex essential for genome organization (Rao *et al*, 2017; Wendt *et al*, 2008)(Fig. S1A). We also derived a regulator-regulator network and focused on the MAX-centered subnetwork using MAX peaks on chromosome 1. The inferred network recovered the canonical MYC-MAX and MXI1-MAX links and further connected the MXI1-MAX branch to the SIN3A-HDAC1 co-repressor module (Tiana *et al*, 2018; Zervos *et al*, 1994)(Fig. S1B). These results demonstrate that ChromBERT-tools effectively leverages regulation-informed embeddings to capture intrinsic interaction potentials between regulators in a context-aware manner.

#### 2.2.2 CRE characterization

Characterizing CREs is fundamental to elucidating the logic of gene expression, as it provides the functional blueprint of the genome. ChromBERT-tools generates cell-type-agnostic embeddings for CREs; pairwise similarity between these embeddings can prioritize functionally related CRE pairs that might reflect shared region-level regulatory programs.

Here, we demonstrate this capability by addressing the classic challenge of CRE characterization. ChromBERT-tools provides the “infer_ep” subcommand as an end-to-end workflow for prioritizing enhancer-promoter (E-P) links, the mechanism by which enable distal CREs to regulate transcription (Fullwood *et al*, 2009). Here we assumed that distal regions exhibited higher embedding similarity with promoter of interest shared the similar regulatory programs with the given promoter, which give us a hint for E-P links. We applied this workflow to ATAC-seq peaks in human embryonic stem cells and prioritized candidate E-P links genome-wide (Supplementary Methods), identifying the *MOB3C*and *RNVU1-15*loci as representative examples. Validation against public RNA Pol II ChIA-PET interaction maps (Consortium, 2012) confirmed that those predicted E-P link from ChromBERT-tools have chromatin contacts (Fig. S1C). This suggests that embeddings from ChromBERT-tools successfully capture the functional features underlying 3D chromatin contact and distal transcriptional regulation.

#### 2.2.4 Cell-type-specific key regulator identification

Cellular identity is governed by distinct regulatory programs orchestrated by a select group of cell-type-specific transcription regulators. ChromBERT-tools identifies these key regulators by fine-tuning ChromBERT using cell-type-specific chromatin accessibility profiles as learning objectives to derive embeddings adapted to the given cell type. Then the tool prioritizes TRs based on the embedding shift, which quantifies each regulator’s divergence in embedding between highly accessible regions and genomic background (low chromatin accessibility) regions. This approach is based on the assumption that key regulators exert the most significant influence on the regulatory program of open chromatin in a given cell type. ChromBERT provides an end-to-end subcommand “infer_cell_key_regulator” to implement this (Supplementary Methods).

We applied this subcommand to B cells, where the model successfully prioritized canonical master regulators, including BCL6, PAX5, POU2F2, ETS1, and MEF2C (Laidlaw and Cyster, 2021). Similarly, in macrophages, the top-ranked candidates included established master regulators such as RUNX1, CEBPA, CEBPB, PPARG, and SPI1 (Fig. S2) (Odegaard *et al*, 2007; Tussiwand and Gautier, 2015). Furthermore, a comparison of fast versus full modes (see Section 2.1.3) revealed that fast mode substantially reduced computation runtime while maintaining comparable performance (Fig. S2).

#### 2.2.5 Cell-state transition driver identification

Cell state transitions are governed by coordinated changes in regulatory programs driven by a subset of key regulators. Identifying the regulators that drive these programs is critical for understanding cell fate control and rationally prioritizing factors for cell-state engineering. ChromBERT-tools fine-tunes model using chromatin accessibility or transcriptome changes between the start and target states to derive transition-informed regulator embeddings. The tool prioritizes candidate drivers via an embedding-shift score that contrast regions gaining activity in the target state against unchanged regions. Larger shifts are assumed to correspond to a regulator’s prominent role in driving the specific state change. This end-to-end workflow is implemented in the “find_driver_in_transition” subcommand (Supplementary Methods).

We applied the workflow to fibroblast-to-iPSC reprogramming, where the top-ranked drivers included the known reprogramming factors POU5F1, SOX2, and NANOG (Schwarz *et al*, 2014; Yu *et al*, 2007). Similarly, in fibroblast-to-myoblast reprogramming, the tool prioritized known myogenic drivers such as MYOD1 (Figs. S3-S4) (Kabadi *et al*, 2015), These results validate the efficacy of transition-specific fine-tuning in isolating the key regulators underlying cell-fate changes. Furthermore, benchmarking in this task revealed that the fast mode also substantially reduced runtime while maintaining prediction performance comparable to the full mode (Fig. S3, S4).

#### 2.2.6 Context-specific cofactor identification

Many transcription regulators, particularly histone modifiers, exhibit functional plasticity: they can bind to distinct classes of genomic loci with different cofactor dependencies, thereby mediating diverse regulatory outcomes within the same cell type (Hu *et al*, 2020). ChromBERT-tools dissects these mechanisms by generating context-specific regulator embeddings via fine-tuning on two user-defined sets of genomic regions and using the class label as learning objective. The workflow prioritizes cofactors as the regulators having high embedding similarity with target regulator in one class of context and have low embedding similarity on another class of context. This end-to-end workflow is implemented in the “find_context_specific_cofactor” subcommand (Supplementary Methods).

We validated this approach using EZH2, a regulator known for its dual roles (Hu *et al*, 2020; Margueron and Reinberg, 2011; Xu *et al*, 2012). The workflow successfully recovered established cofactors associated with its canonical repressive function (H3K27me3-dependent). Crucially, it also highlighted candidate cofactors for its non-canonical functions (H3K27me3-independent), including E2F1 and STAT3. These findings align with recent reports on the context-dependent, non-repressive roles of EZH2 (Hu *et al*, 2020) (Fig. S5).

### 2.3 Cistrome imputation

Despite the availability of numerous ChIP-seq datasets, TR occupancy profiles remain incomplete across the vast majority of cell types. ChromBERT-tools generates regulator and region embeddings that provide regulator and region information. When combined with cell-type-specific chromatin accessibility information, these representations enable imputation of cell-type-specific TF occupancy at candidate regulatory regions. To facilitate this, ChromBERT-tools includes the “impute_cistrome” subcommand, an end-to-end workflow for genome-wide occupancy prediction of TRs in the given cell type (Supplementary Methods).

We demonstrated this capability by imputing cistromes of BRD4 in several cell types. Evaluated against corresponding BRD4 ChIP-seq in K562 and A549 cells, the imputation achieved AUPRCs of 0.66 and 0.54 (outperforming baseline at 0.36 and 0.47, respectively). Representative genome-browser views show concordant predicted and experimental BRD4 tracks at selected loci (Fig. S6), supporting the effectiveness and accuracy of cistrome imputation.

### 2.4 Implementation

ChromBERT-tools is implemented in Python (v3.9+) and distributed with a containerized environment (Apptainer/Singularity) to simplify installation and promote consistent deployment across systems. The source code and documentation, including command-line options, input/output specifications, and tutorials, are publicly available (https://github.com/TongjiZhanglab/ChromBERT-tools; https://chrombert-tools.readthedocs.io/en/latest/).

## 3 Conclusion

We developed ChromBERT-tools, an end-to-end toolkit with unified CLIs and Python APIs that transforms a technically demanding regulatory foundation model into a practical resource for generating, reusing, and interpreting regulation-informed embeddings. ChromBERT-tools integrates and automates practical steps, including multi-genome/multi-resolution configuration, standardized raw-input ingestion, task-specific fine-tuning configuration, built-in run validation and downstream biological interpretation of embeddings.

ChromBERT-tools enables both cell-type-agnostic and cell-type-specific regulator embeddings and supports core applications such as key regulator identification, transition driver prioritization, and cistrome imputation. ChromBERT-tools lowers the engineering barrier to applying regulatory foundation models for routine regulatory inference and candidate prioritization across diverse biological contexts, and provides a modular foundation for incorporating new datasets, multi-modal inputs, additional model backends, and downstream workflows.

## Acknowledgements

We thank Dongxu Yang for comments on the design, and the authors of ChromBERT for support and feedback.

## Author contributions

Qianqian Chen (Formal analysis [lead], Investigation [lead], Methodology [lead], Software [lead], Validation [lead], Visualisation [lead], Writing—original draft [equal], Writing—review & editing [equal]), Zhaowei Yu (Conceptualization [lead], Project administration [lead], Resources [lead], Supervision [lead], Funding acquisition [lead], Writing—original draft [equal], Writing—review & editing [lead]), Yong Zhang (Conceptualization [lead], Project administration [lead], Resources [lead], Supervision [lead], Funding acquisition [lead], Writing—original draft [equal], Writing—review & editing [lead]).

