## Supplementary Materials for "ChromBERT-tools: A versatile toolkit for context-specific embedding of transcription regulators across different cell types"

### Supplementary figure

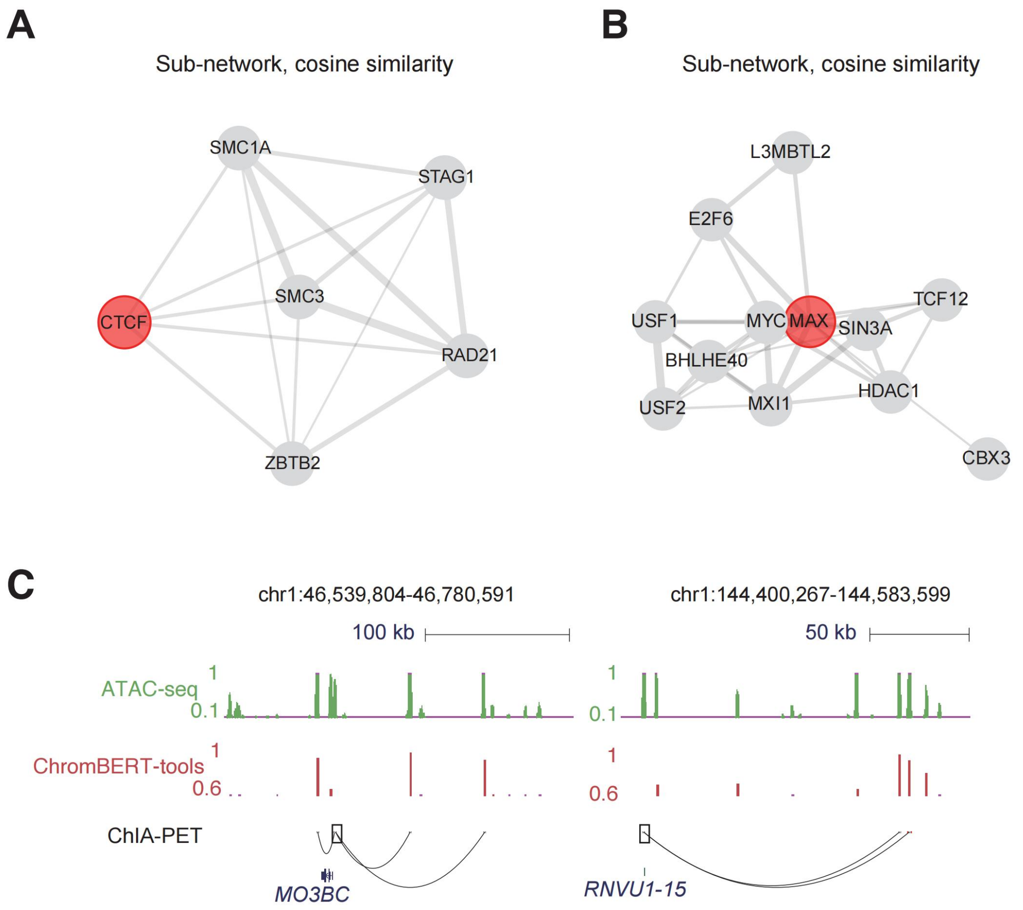


Figure S1. Interpretation of cell-type-agnostic regulation-informed embeddings.

(A-B) Regulator-regulator similarity subnetworks inferred from cell-type-agnostic regulator embeddings using the regions of interest; edge opacity reflects pairwise embedding similarity, and only edges with similarity above the 98th percentile of all pairwise similarities were retained.

(A) CTCF-centered subnetwork using CTCF peaks on chromosome 1 (ENCODE: ENCFF664UGR[1]).

(B) MAX-centered subnetwork using MAX peaks on chromosome 1 (ENCODE: ENCFF266YHW[1]).

(C) UCSC Genome Browser views showing representative enhancer-promoter contacts for MOB3C (chr1:46,539,804-46,780,591; left) and RNVU1-15 (chr1:144,400,267-144,583,599; right) and their embedding similarities. Tracks from top to bottom: ATAC-seq signal in hESCs (GSM2386582[2]; green), cosine similarity between general (pre-trained) region embedding vectors for TSS-located 1kb bin (boxed) and nearby accessible distal 1kb-bins within ±250 kb (red), and chromatin loops detected by Pol II ChIA-PET in hESCs (ENCSR782EKZ[1]; blue arcs). Only loops linking accessible distal regions to the target promoters are shown.

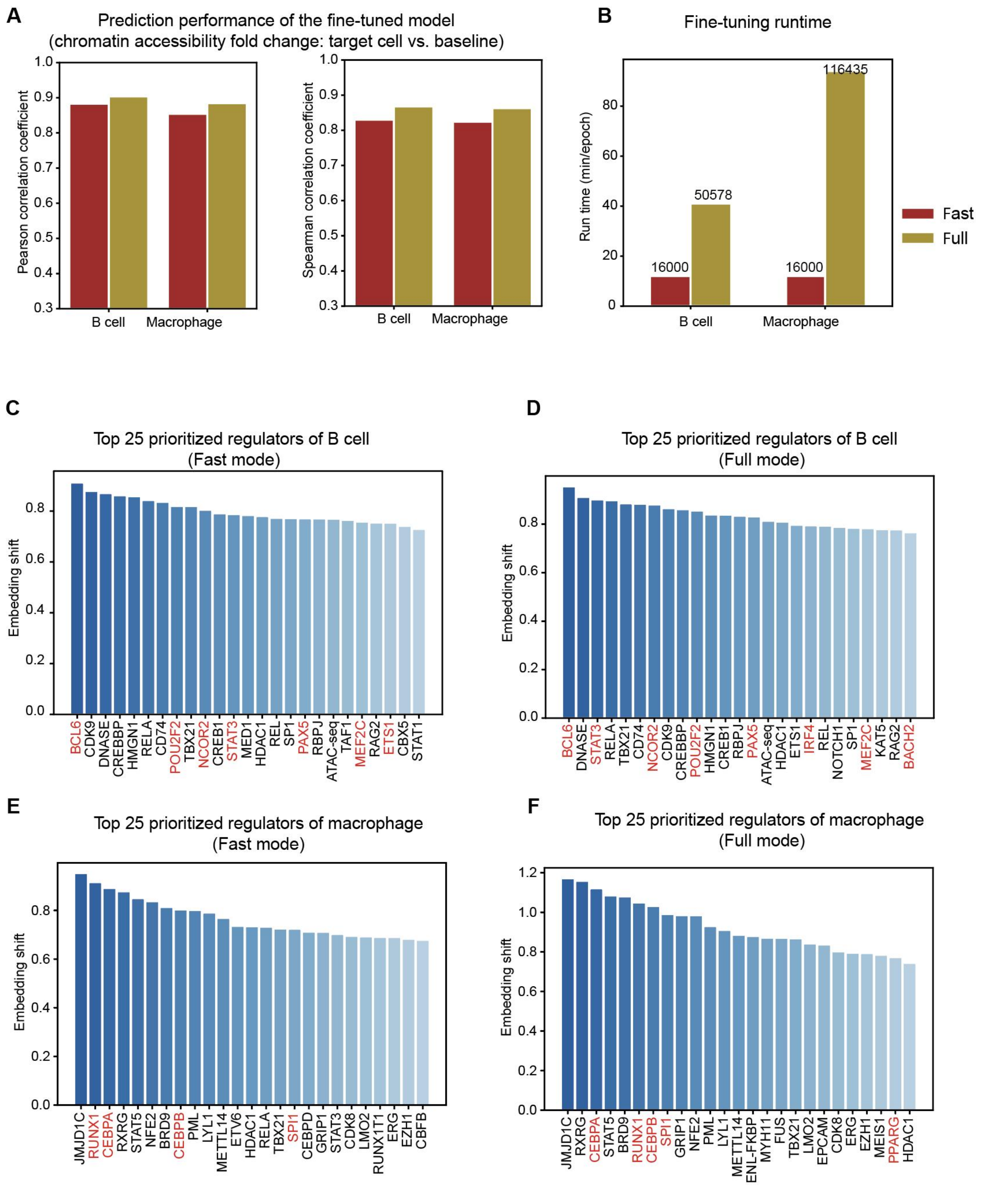


Figure S2. Interpretation of cell-type-specific regulation-informed embeddings: key regulator identification.

(A) Bar plots comparing fast and full modes for predicting genome-wide chromatin accessibility changes from a baseline reference to a target cell type. As a baseline, ChromBERT-tools provides a precomputed mean accessibility signal for each reference bin, aggregated across all available accessibility cistromes in the metadata; see details in the Supplementary Methods. DNase-seq datasets (ENCODE): B cell (ENCSR468AKF[1]) and macrophage (ENCSR593WXK[1]).

(B) Bar plots comparing fast and full modes runtimes for fine-tuning. Numbers on the bars indicate the number of training regions.

(C-F) Top 25 regulators prioritized in B cells and macrophages using cell-type-specific (fine-tuned) regulator embeddings (fast and full modes). Bar height indicates the embedding-shift score, defined as the difference between each regulator's embedding aggregated over high-accessibility regions and over low-accessibility background regions within the same cell type (see Supplementary Methods). Known lineage master regulators are highlighted in red.
(C) B cells, fast mode. (D) B cells, full mode. (E) Macrophages, fast mode. (F) Macrophages, full mode.

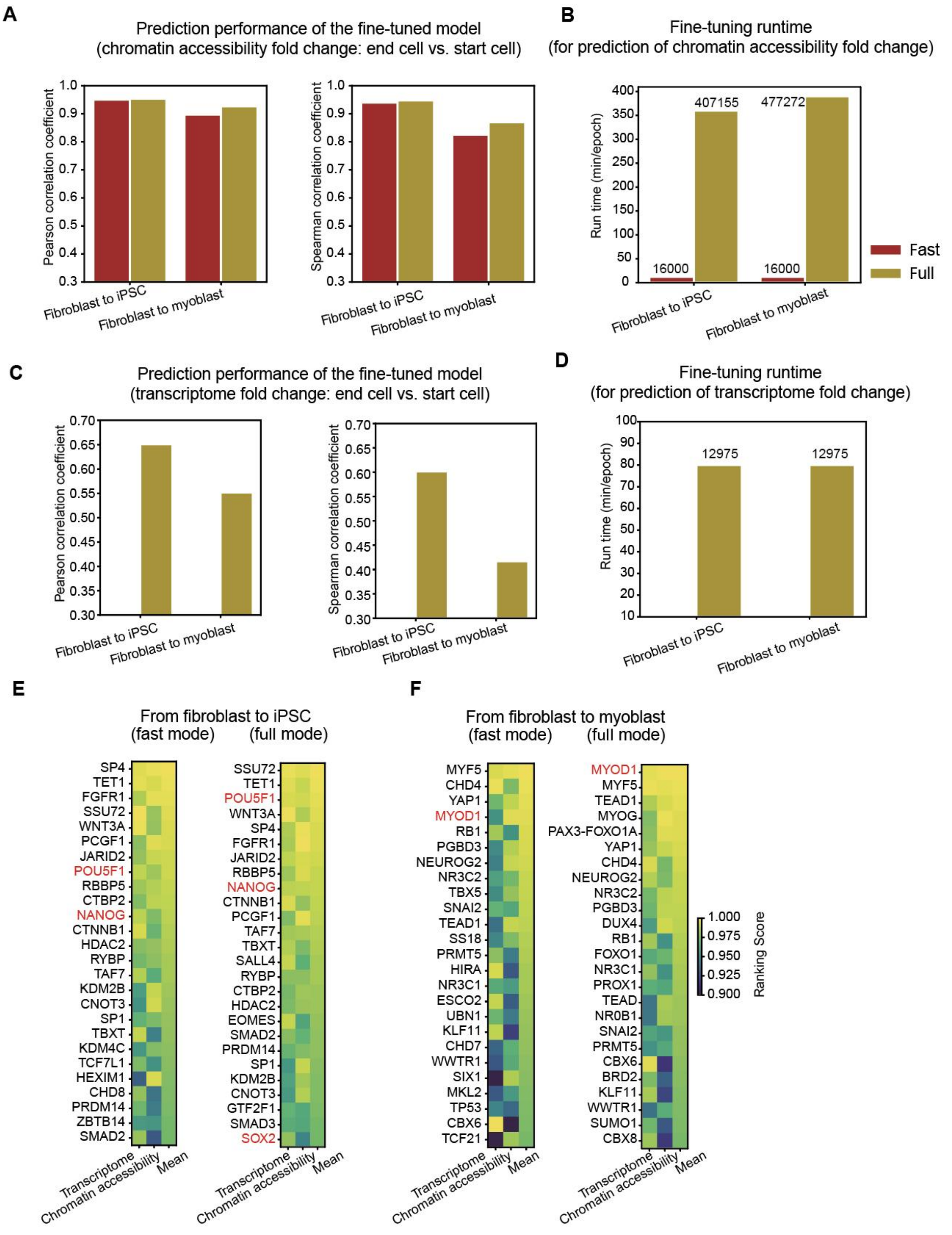


Figure S3. Interpretation of cell-type-specific regulation-informed embeddings: key driver identification for cell-state transitions.

(A) Bar plots comparing fast mode and full mode in predicting genome-wide chromatin accessibility changes during cell-state transitions. DNase-seq datasets (ENCODE): fibroblast (ENCFF184KAM[3]), iPSC (ENCFF540VPT[3]), and myoblast (ENCFF647RNC[3]).

(B) Bar plots comparing fine-tuning runtime between fast and full modes for the task in (A). Numbers on the bars indicate the number of training regions.

(C) Bar plots showing full-mode performance for predicting genome-wide transcriptome changes during cell-state transitions. Because expression prediction is restricted to protein-coding genes (< 20,000 genes), we report full-mode results. Transcriptome profiles derived from CAGE-seq in seven cell types available in the FANTOM5 database[4].
(D) Bar plots showing fine-tuning runtime for the task in (C). Numbers on the bars indicate the number of training regions.
(E) Heatmap of ranking scores for the top 25 regulators prioritized in the fibroblast-to-iPSC transition (fast mode: left; full mode: right). Known iPSC reprogramming factors are annotated and highlighted in red.
(F) Heatmap of ranking scores for the top 25 regulators identified in the fibroblast-to-myoblast transition (fast mode: left; full mode: right). Known myoblast reprogramming factors are highlighted in red.

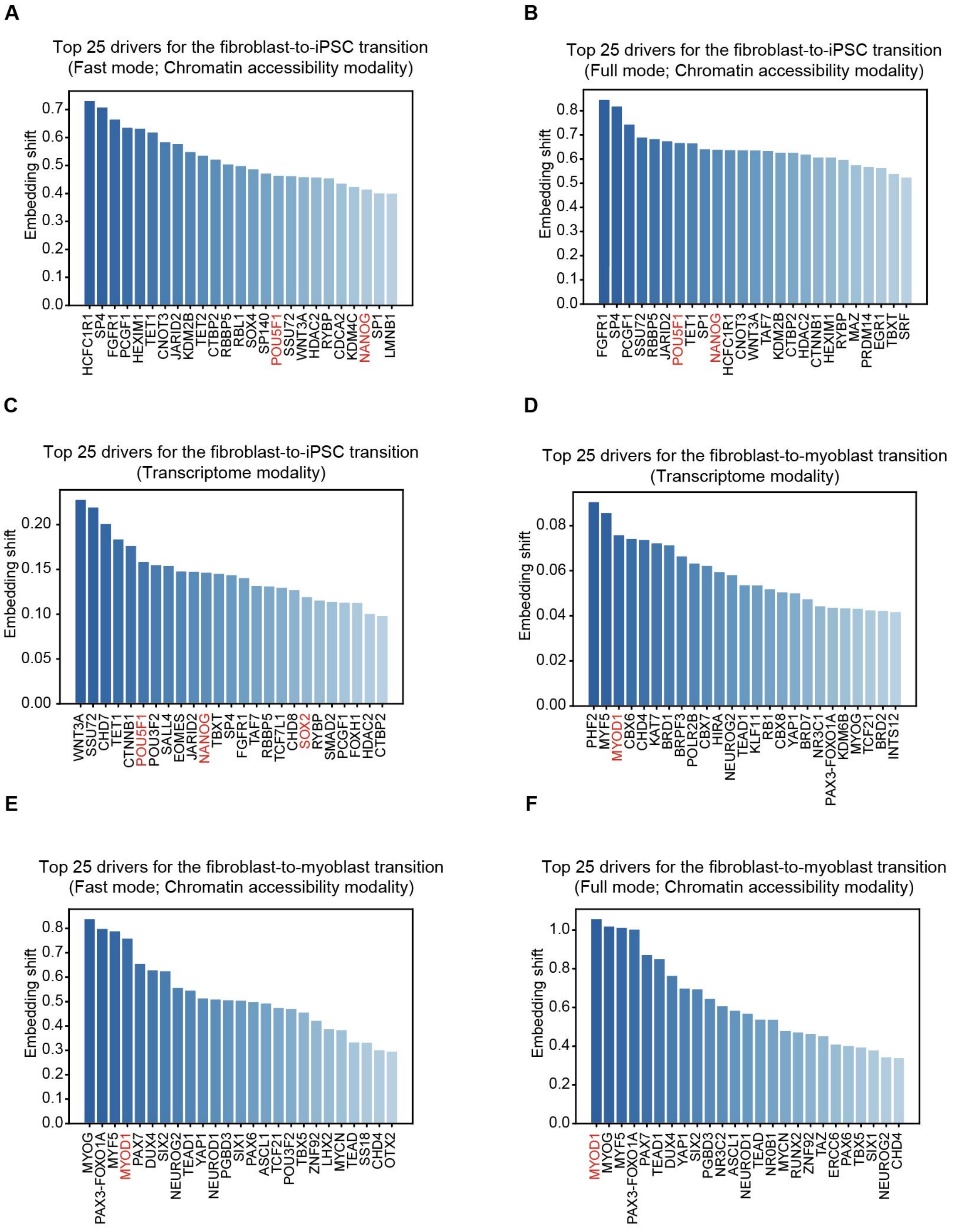


Figure S4. Interpretation of cell-type-specific regulation-informed embeddings: key driver identification for cell-state transitions using a single modality.

(A-F) Top 25 regulators prioritized for fibroblast-to-iPSC and fibroblast-to-myoblast transitions using cell-specific (fine-tuned) regulator embeddings derived from a single modality (chromatin accessibility or transcriptome), comparing fast and full fine-tuning modes where applicable. Bar height indicates the embedding-shift score, defined as the distance between each regulator's embedding for gained-activity regions and for unchanged regions (see Supplementary Methods). Known reprogramming factors are highlighted in red. DNase-seq datasets (ENCODE): fibroblast (ENCFF184KAM[3]), iPSC (ENCFF540VPT[3]), and myoblast (ENCFF647RNC[3]). Transcriptome profiles derived from CAGE-seq in seven cell types available in the FANTOM5 database[4].

(A) Fibroblast-to-iPSC, chromatin accessibility modality, fast mode.

(B) Fibroblast-to-iPSC, chromatin accessibility modality, full mode.

(C) Fibroblast-to-iPSC, transcriptome modality, full mode only (< 20,000 protein-coding genes).

(D) Fibroblast-to-myoblast, transcriptome modality, full mode only (< 20,000 protein-coding genes).

(E) Fibroblast-to-myoblast, chromatin accessibility modality, fast mode.

(F) Fibroblast-to-myoblast, chromatin accessibility modality, full mode.

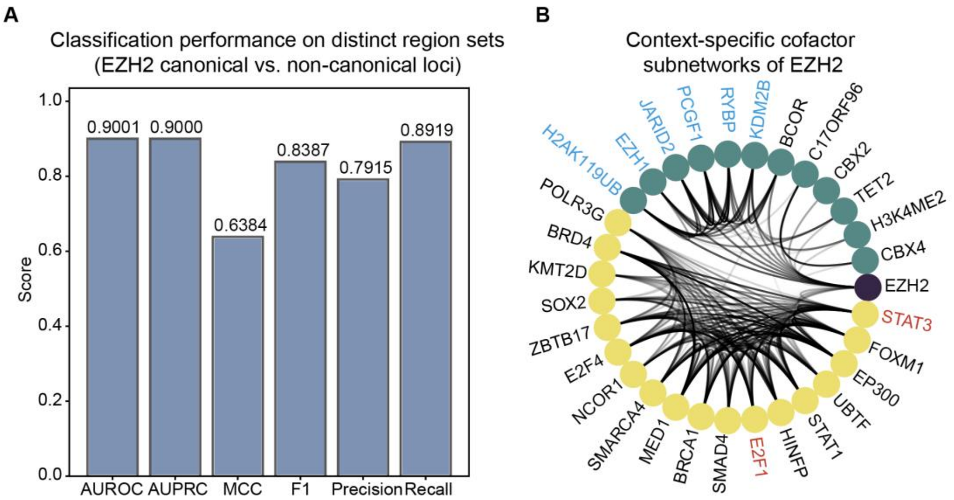


Figure S5. Interpretation of cell-type-specific regulation-informed embeddings: context-specific cofactor identification.

(A) Bar plots showing classification performance for EZH2 canonical versus non-canonical loci. Canonical loci are EZH2-bound regions with H3K27me3, whereas non-canonical loci are EZH2-bound regions without H3K27me3. To mitigate H3K27me3-driven effects, ChromBERT-tools excludes H3K27me3-associated cistromes from the reference inputs for this task. ChIP-seq data (GEO): EZH2 (GSE29611[1]) and H3K27me3 (GSE61176[5]).

(B) Circos plot of EZH2 embedding similarities at canonical (H3K27me3+) and non-canonical (H3K27me3−) loci, highlighting two regulator groups with locus-preferential similarity to EZH2 (see Supplementary Methods). Regulators were assigned to the canonical group (green) if their EZH2 similarity at canonical loci ranked in the top 5% across regulators and exceeded that at non-canonical loci by > 0.1; the non-canonical group (yellow) showed the converse pattern. Known canonical EZH2 cofactors are highlighted in blue, and reported non-canonical EZH2 cofactors (E2F1 and STAT3) are highlighted in red. Nodes represent regulators and edge opacity indicates regulator–regulator embedding similarity.

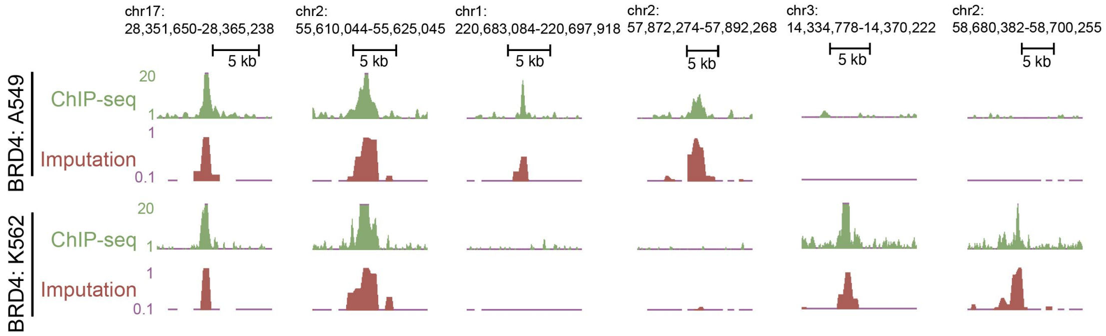


Figure S6. Cistrome imputation. BRD4 cistrome imputation in A549 and K562 at the indicated locus using cell-type-agnostic BRD4 and region embeddings, together with DNase-derived cistrome embeddings for each cell type (K562 DNase-seq: GEO: GSM623516[6]; A549 DNase-seq: GEO: GSM1014517[7]). Imputed signal is shown in red and measured BRD4 ChIP-seq (ground truth) in green (K562 BRD4 ChIP-seq: GEO: GSM2635249[8]; A549 BRD4 ChIP-seq: GEO: GSM2359441[9]).

#### Supplementary table

| Application | CLI & API | Input | Output | Fine-tune |
|---|---|---|---|---|
| Generation of cell-type-agnostic embeddings | embed_gene | ➢ Gene Ensembl ID or gene symbol | Gene embedding | No |
| | embed_region | ➢ Regions (BED) | Region embedding | |
| | embed_cistrome | ➢ Cistrome identifier (e.g., factor:cell);<br>➢ Regions (BED) | Cistrome embedding | |
| | embed_regulator | ➢ Regulators name;<br>➢ Regions (BED) | Regulator embedding | |
| Regulator-regulator interaction | infer_regulator_network | ➢ Regions (BED);<br>➢ (optional)Regulator name | Regulator-regulator edge list; Subnetwork of regulator | No |
| Cistrome imputation | impute_cistrome | ➢ Cistrome identifier (e.g., factor:cell);<br>➢ Regions (BED) | Per-region binding probability | No |
| CRE characterization (enhancer-promoter loop) | infer_ep | ➢ Regions (BED) | Enhancer-promoter region similarity | No |
| Generation of cell-type-specific embeddings | embed_cell_gene | ➢ Gene ID/symbol;<br>➢ (optional) peaks (BED) + signal (bigWig);<br>➢ (optional)fine-tuned checkpoint | Cell-type-specific gene embedding | Optional |
| | embed_cell_cistrome | ➢ Gene ID/symbol;<br>➢ (optional) peaks (BED) + signal (bigWig);<br>➢ (optional)fine-tuned checkpoint | Cell-type-specific cistrome embedding | |
| | embed_cell_regulator | ➢ Regulator name;<br>➢ Regions (BED);<br>➢ (optional) peaks (BED) + signal (bigWig);<br>➢ (optional) checkpoint | Cell-type-specific regulator embedding | |
| | embed_cell_region | ➢ Regions (BED);<br>➢ (optional) peaks+signal;<br>➢ (optional) checkpoint | Cell-type-specific region embeddings | |
| Cell-type-specifi | infer_cell_key_regulat | ➢ Accessibility peaks | Cell-type-speci | Optio |

| | | | | |
|---|---|---|---|---|
| c key regulator identification | or | (BED) + signal track (bigWig);<br>➢ (optional) fine-tuned checkpoint | fic regulator ranking (csv) | nal |
| Cell-state dirver identification | find_driver_in_transition | ➢ Start and end cell-state accessibility accessibility (peaks BED + signal bigWig);<br>➢ (optional)RNA-seq expression (TPM) ;<br>➢ (optional) checkpoint | Regulator importance ranking (csv) | Optional |
| Context-specific cofactor identification | find_context_specific_cofactor | ➢ Region set A (BED) + region set B (BED);<br>➢ (optional) checkpoint;<br>➢ (optional) the dual function of regulators;<br>➢ (optional) checkpoint | Regulator importance ranking (csv);<br>(optional) Dual-functional regulator subnetwork | Optional |

Note:
"Optional" indicates that fine-tuning can be skipped if a fine-tuned checkpoint is provided; "No" indicates that no fine-tuning is involved;
BED: standard three-column genomic intervals (chrom, start, end);
bigWig: genome-wide signal track;
TPM: Transcripts Per Million;
CLI: Command-line interfaces
Detailed in: https://chrombert-tools.readthedocs.io/en/latest/

**Table S1. Summary of ChromBERT-tools interfaces and applications**

### Supplementary methods

#### Data processing

**Region-to-bin mapping.** ChromBERT-tools takes the first three columns (chrom/start/end) from user-provided BED files, sorts by genomic coordinates, and maps them to ChromBERT reference bins using *bedtools[10] intersect -a <ref_bins> -b <user_regions> -f 0.5 -F 0.5 -wa -wb -e*; for each hit ChromBERT-tools outputs the user region coordinates and the matched reference-bin identifier (bin_id) (one-to-many mapping allowed).

**Gene-to-bin mapping.** ChromBERT-tools distributes a precomputed gene-to-bin mapping table, in which each gene is assigned to the reference bin containing its transcription start site (TSS) as defined from the GRCh38 Ensembl GTF (release 110). Given a gene symbol or Ensembl gene ID, ChromBERT-tools looks up the corresponding reference-bin identifier using this table.

**Regulators and cistromes**. ChromBERT-tools provides curated regulator lists (human: 1,073; mouse: 703) and a metadata table listing all available cistromes (cell type-factor pairs). Given a regulator name or cistrome name, ChromBERT-tools resolves the input against these resources (e.g., for lookup and validation).

**BigWig signal summarization.** ChromBERT-tools summarizes signals from user-provided ATAC-seq/DNase-seq BigWig tracks over mapped intervals derived from user-provided regions using pybbi.stackup[11] (default: mean over the interval); optionally, values are normalized by the BigWig's global mean.

**Cell-type-specific supervision signal for fine-tuning**. For cell-type-specific key regulator identification, ChromBERT-tools constructs fine-tuning inputs from user-provided raw chromatin accessibility data (peak BED and BigWig tracks). Using the region-to-bin mapping and BigWig signal summarization steps, ChromBERT-tools computes the mean accessibility signal $s_{c,b}$ for each reference bin b in the target cell type c. As a baseline, ChromBERT-tools provides a precomputed mean accessibility signal $s_{baseline,b}$ for each reference bin, aggregated across all available accessibility cistromes in the metadata. ChromBERT-tools apply a log transform to both signals and define the supervised training target (fold-change) as:

$$y_b = log_2(1 + s_{c,b}) - log_2(1 + s_{baseline,b})$$

ChromBERT-tools defines highly accessible regions as reference bins with $y_b$ > 1 and select the top 1,000 bins ranked by $y_b$, and ChromBERT-tools defines background regions by first filtering bins with $s_{c,b}$ > 0 and $s_{baseline,b}$ > 0 and and then selecting the 1,000 bins with the smallest $|y_b|$.

**Cell-state transition supervision signal for fine-tuning**. For transition-specific key regulator identification, ChromBERT-tools constructs fine-tuning inputs from user-provided chromatin accessibility data of the start and end cell states (peak BED and BigWig tracks). ChromBERT-tools defines training regions as the union of peaks from both states together with all protein-coding gene TSS bins. For each reference bin b, ChromBERT-tools computes the mean accessibility signal in the start and end states, denoted $s_{start,b}$ and $s_{end,b}$.The supervision target is the log-transformed accessibility change:

$$y_b = log_2(1 + s_{end,b}) - log_2(1 + s_{start,b})$$

If RNA expression data are provided, ChromBERT-tools adds a gene-level supervision target for each gene g and predicts protein-coding genes only:

$$y_g = ln(1 + e_{end,g}) - ln(1 + e_{start,g})$$

where $e_{*,g}$ denotes the expression level of gene g (e.g., TPM).

ChromBERT-tools defines up regions as reference bins with $y_b$ > 1 and select the top 1,000 bins ranked by $y_b$. No-change bins are defined by requiring $s_{end,b}$ > 0 and $s_{start,b}$ > 0, and then selecting the 1,000 bins with the smallest $|y_b|$. Similarly, up genes are defined as genes with $y_g$ > 1, and no-change genes are selected as the 1,000 genes with the smallest $|y_g|$ (optionally restricting to $|y_g|$ < 0.5 to exclude large expression changes).

**Binary supervision for distinct region sets.** For key regulator identification from distinct region sets, ChromBERT-tools constructs fine-tuning inputs from two user-defined region sets (A and B). Regions in set A are labeled as positive (1/True) and regions in set B as negative (0/False), forming a binary classification task.

All fine-tuning datasets are split into training/validation/test sets with an 8:1:1 ratio.

#### Model fine-tuning

For classification tasks, ChromBERT-tools optimizes the model using binary cross-entropy (BCE) loss and reports the area under the precision-recall curve (AUPRC) on the validation set. For regression tasks, ChromBERT-tools uses root mean squared error (RMSE) loss and reports the Pearson correlation coefficient (PCC) between predictions and targets on the validation set. ChromBERT-tools applies early stopping when the monitored validation metric does not improve for 5 consecutive validation checks (min_delta = 0.01), except for expression prediction, where 10 consecutive validation checks. ChromBERT-tools selects and saves the checkpoint with the best validation performance (highest AUPRC/PCC, depending on the task). Training runs for up to 10 epochs. Gradients are accumulated over 64 mini-batches (accumulate_grad_batches = 64). Validation is performed five times per epoch (every 20% of an epoch). By default, ChromBERT-tools freezes the first six Transformer blocks and fine-tunes only the final two blocks and the task-specific head.

#### Model validation

ChromBERT-tools evaluates the fine-tuned model on a held-out test set, reporting PCC for regression tasks and AUPRC for classification tasks. To mitigate occasional training instability due to random initialization and local optima, if the test metric falls below 0.2, ChromBERT-tools automatically restarts fine-tuning with a new initialization; this is attempted up to three times. ChromBERT-tools also reports additional test-set metrics (Classification: AUROC, MCC, F1, Precision, Recall; Regression: Spearman correlation coefficient, MSE, MAE).

#### Embedding analysis

ChromBERT-tools computes embedding similarity and embedding-shift scores as described in the main text (Embedding analysis); additional task-specific analysis steps are applied when needed.

**For regulator-regulator interaction analysis (subcommand: infer_regulator_network)**, ChromBERT-tools defines an edge between a pair of regulators if their embedding cosine similarity exceeds a user-defined threshold (default: the 98th percentile of all regulator-regulator cosine similarities), indicating a putative interaction. For regulator-subnetwork visualization, ChromBERT-tools extracts the k-hop neighborhood of a user-specified regulator in the regulator-regulator graph (default: k=1). These similarities are computed using the cell-type-agnostic embeddings without task-specific fine-tuning; regulator embeddings are obtained by averaging across user-provided regions.

**For CRE characterization (enhancer-promoter loop) analysis (subcommand: infer_ep)**, ChromBERT-tools quantifies each enhancer regions (defined open regions by user-provided ATAC-seq or DNase-seq) with each gene TSS region bins embedding cosine similarity, their distance shorted +- 250 kb, with the higher embedding cosine similarity, means there are more probabilities exist enhancer-promoter loops. These similarities are computed using the cell-type-agnostic embeddings without task-specific fine-tuning.

**For cistrome imputation (subcommand: impute_cistrome)**, ChromBERT-tools extracts cell-type-agnostic regulator embeddings and cell-type accessibility embeddings from user-provided cistrome names, and extracts region embeddings for user-provided genomic regions. ChromBERT-tools performs cistrome imputation using the task-specific head and weights distributed with the original ChromBERT release, requiring no further fine-tuning, it predicts regulator binding probabilities across the input regions for the specified cell type. For comparison, we also provide a baseline method, which assigns each region a score equal to the cell-type DNase-seq signal multiplied by an indicator of whether the region overlaps a regulator-specific candidate peak set (defined from ≥ 5 related reference cistromes for the same regulator (across other cell types) or motif-based bins when fewer are available).

**For cell-specific key regulator identification (subcommand: infer_cell_key_regulator)**, ChromBERT-tools quantifies each regulator’s embedding shift between the highly accessible regions and background region sets (see Data processing above) by comparing the mean cell-type-specific regulator embeddings computed over each set and reports the top 25 regulators with the largest shift scores as candidate key regulators.

**For cell-state transition driver identification (subcommand: find_driver_in_transition)**, ChromBERT-tools computes each regulator’s mean embedding on the up and no-change sets (see Data processing above) separately for chromatin accessibility (region sets) and gene expression (gene sets). It then quantifies the embedding shift between the two means and reports the top 25 regulators with the largest shift scores for each modality. For multimodal ranking, ChromBERT-tools combines the two modalities by averaging each regulator’s rank across accessibility and expression.

**For context-specific cofactor identification (subcommand: find_context_specific_cofactor)**, ChromBERT-tools computes each regulator’s mean embedding on the positive and negative region sets (see Data processing above). It then quantifies the embedding shift between the two means and reports the top 25 regulators with the largest shift scores. For user-specified dual-function regulators (e.g., EZH2 in our case study), ChromBERT-tools additionally identifies distinct co-factors by computing cosine similarity between the specified regulator and all other regulators on the positive and negative region sets. Regulators with similarity above a user-defined threshold (default: 95th percentile) are considered candidate co-factors. A co-factor is labeled positive if it is > 0.1 more similar on the positive set than on the negative set, and negative if the reverse holds (> 0.1).
